# Co-Occurrence Network Analysis Reveals The Alterations Of The Skin Microbiome And Metabolome In Atopic Dermatitis Patients

**DOI:** 10.1101/2023.08.17.553735

**Authors:** Paulo Wender P. Gomes, Helena Mannochio-Russo, Junhong Mao, Haoqi Nina Zhao, Craig D. Tipton, Jacob Ancira, Pieter C. Dorrestein, Min Li

**Author notes:** **Corresponding author’s email address** Address correspondence to.

## Abstract

Skin microbiome can be altered in patients with Atopic Dermatitis (AD). An understanding of the changes from healthy to atopic skin can help develop new targets for better treatments and identify specific microbial or molecular biomarkers. This study investigates the skin microbiome and metabolome of healthy subjects and lesion (ADL) and non-lesion (ADNL) of AD patients by 16S rRNA gene sequencing and mass spectrometry, respectively. Samples from AD patients showed alterations in the diversity and composition of the skin microbiome. *Staphylococcus* species, especially *S. aureus*, were significantly increased in the ADL group. Metabolomic profiles were also different between the groups. Dipeptide-derived are more abundant in ADL, which may be related to skin inflammation. Co-occurrence network analysis was applied to integrate the microbiome and metabolomics data and revealed higher co-occurrence of metabolites and bacteria in healthy and ADNL compared to ADL. *S. aureus* co-occurred with dipeptide-derived in ADL, while phytosphingosine-derived compounds showed co-occurrences with commensal bacteria, *e*.*g. Paracoccus* sp., *Pseudomonas* sp., *Prevotella bivia, Lactobacillus iners, Anaerococcus* sp., *Micrococcus* sp., *Corynebacterium ureicelerivorans, Corynebacterium massiliense, Streptococcus thermophilus*, and *Roseomonas mucosa*, in healthy and ADNL groups. Therefore, these findings provide valuable insights into how AD affects the human skin metabolome and microbiome.

**Importance:** This study provides valuable insight into changes in the skin microbiome and associated metabolomic profiles. It also identifies new therapeutic targets that may be useful for developing personalized treatments for individuals with atopic dermatitis based on their unique skin microbiome and metabolic profiles.

## Introduction

As the largest organ of the human body, the skin plays a vital role in maintaining a stable internal environment and protecting the body from external factors. The skin’s outer layer is composed of lipids and proteins, as well as skin appendages like hair follicles and eccrine glands, which produce lipids, antimicrobial peptides, enzymes, and salts (1). When the skin is stressed, such as in dermatitis, changes may occur in the microbial community living on the skin and the molecules produced by skin cells.

It is well-established that skin hosts microbes that are essential in the systemic, pathophysiologic, and biochemical balance of the organism (2). These interactions are critical for health and are commonly noticed in the human skin (3). Dermatitis and other stress on the skin can change the molecules derived from the host microbes and disrupt the healthy balance between the host and skin microbiome. Also, inflammatory conditions significantly impact the skin barrier function, resulting in enhanced permeability and water loss (4). This disruption has the potential to modify the skin’s chemical composition by perturbing the levels of essential lipids, including ceramides, cholesterol, and free fatty acids, which play a critical role in maintaining the integrity of the barrier (4). Furthermore, inflammatory processes give rise to reactive oxygen species that can exert an influence on the chemical composition by inducing damage to cellular structures, lipids, and proteins (5). It should be noted that since inflammatory conditions disturb the microbiome, it can also lead to dysbiosis. Dysbiosis refers to an alteration in the composition of microbial communities, which can have significant implications for skin health and its function (6). For instance, conditions such as atopic dermatitis have been closely associated with a reduction in microbial diversity and an overgrowth of specific pathogenic bacteria, such as *S. aureus* (7, 8).

Since the last decade, advances in mass spectrometry and microbial sequencing technologies have allowed us to disclose many microbial groups living in the human skin, which also affects the molecules that can be found in the skin (9–11). In this context, mass spectrometry-based metabolomics and 16S ribosomal RNA (rRNA) gene sequencing have gained increasing attention in the study of skin (12, 13). Metabolomics involves the study of small molecules produced by cells, tissues, or organisms, and how they change in response to various conditions or stress (14–16). On the other hand, microbial sequencing involves the analysis of genetic material (17–19). This approach allows studying the collective genomes of entire communities of microorganisms rather than focusing on individual organisms. Thus, combining these two approaches has proven helpful in identifying changes in the metabolic and microbiome profile of several biological samples (19, 20). Therefore, in this study, we used metabolomics and 16S rRNA gene sequencing approaches to investigate the changes in the metabolome and microbiome of skin swabs of healthy patients and subjects with atopic dermatitis (FIG 1). Furthermore, the two omics data were integrated by co-occurrence network analysis to explore the relationship of metabolites and bacteria in relation to atopic dermatitis.

**FIG 1.**
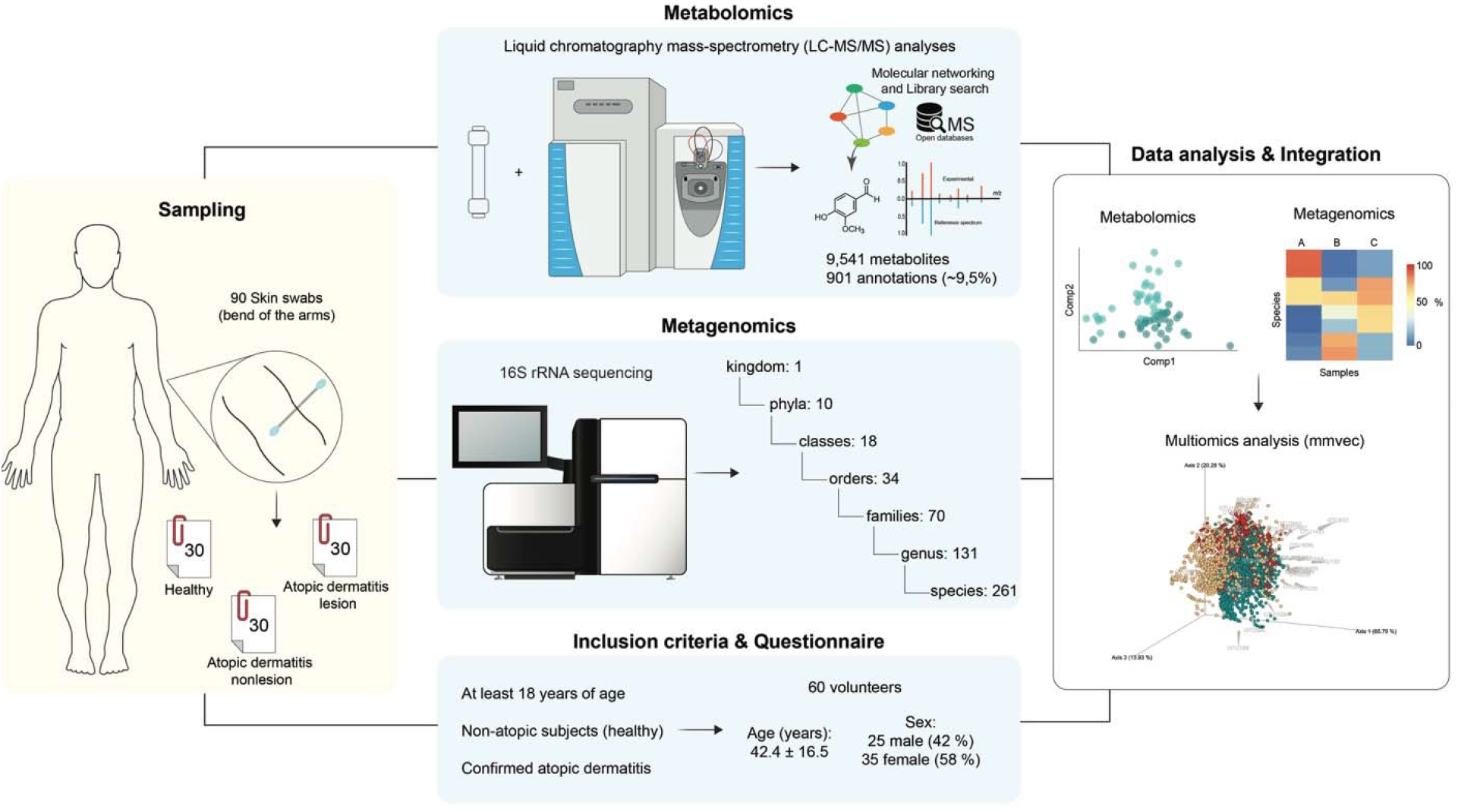
Experimental design and analytical workflow. A total of 60 volunteers were selected based on the inclusion criteria and were included as the subjects of this study. Subsequently, 90 skin swabs were collected from the skin bend of their arms and were divided into three groups: healthy, atopic dermatitis non-lesion (ADNL), and atopic dermatitis lesion (ADL), with 30 swab samples for each group. Each swab sample was analyzed by 16S rRNA sequencing and by liquid chromatography-mass spectrometry. The metabolomics data was analyzed by molecular networking approach and library search. A multi-omic microbe-molecule co-occurrence analysis (mmvec) was used to combine both datasets.

## Results

Skin swabs were collected from healthy subjects (N=30) and lesional and non-lesional (5 cm away from the lesions) skin of the patients with atopic dermatitis (N=30) to gain a deeper understanding of the underlying mechanisms of atopic dermatitis.

### Microbiome profiles by 16S rRNA sequencing

Atopic dermatitis significantly changed the skin microbial composition. The alpha diversity (Shannon diversity index) of AD lesion was significantly lower than AD non-lesion (p<0.01) and healthy group (p=0.01) (FIG 2a), while the diversity of AD non-lesion and healthy groups were not significantly different. The overall microbial profiles of each group at the species level were illustrated in FIG 2b. The most abundant bacterial lineages observed included *Cutibacterium acnes, Staphylococcus sp/epidermidis/aureus*, and *Corynebacterium tuberculostearicum*, accounting for 59% of all read counts study-wide. Beta diversity was assessed using weighted UniFrac distance to summarize the microbial composition between the groups. Using PCoA for qualitative clustering and PERMANOVA for significance testing, AD lesion samples are significantly separated from AD non-lesion and healthy samples (FIG 2c). Then, ANCOM-BC was used to identify six species-level clusters which significantly differed between groups (Table S1). Of these, two species (*S. aureus* and *S. epidermidis*) were much more abundant than the rest and found at significantly greater proportions in AD samples (FIG 2d).

**FIG 2.**
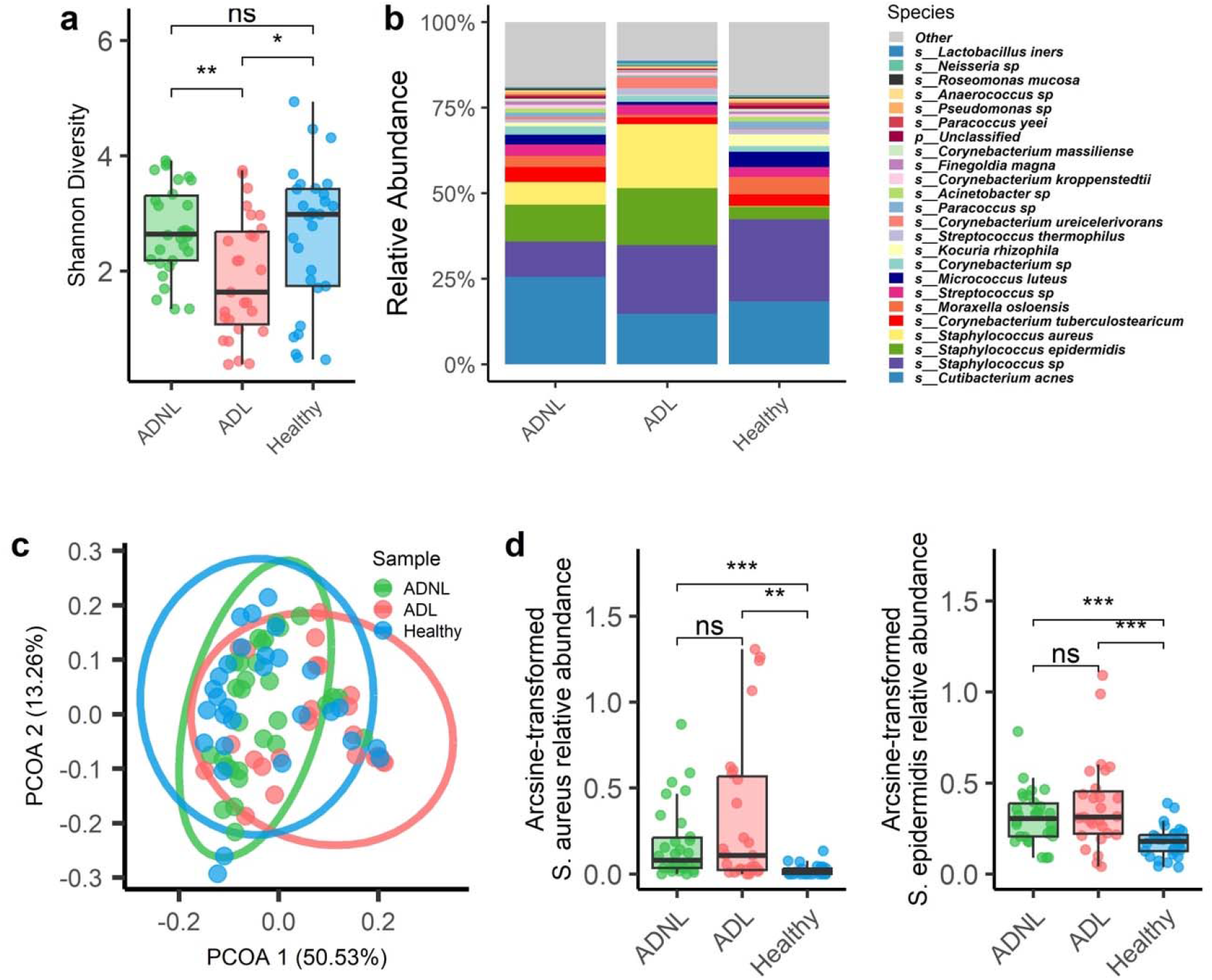
The skin microbiome profiles of lesion and non-lesion of atopic dermatitis patients and healthy subjects characterized by 16S rRNA gene sequencing. a) Shannon diversity of 16S rRNA sequencing, grouped and colored by the three sample groups: health, ADNL, and ADL. Boxplots show the median (designated by the horizontal line), first and third quartiles of each of the three groups. Additionally, ANOVA and Tukey testing was conducted and showed samples to significantly differ between groups, as well as ADL arguing the most between the 2 other sample groups. b) Relative abundance of the top 30 species in each group. Values on the y-axis are percentages (relative abundance) based on the reading count. c) Principal Coordinates Analysis plot based on weighted Unifrac distances of 16S microbiome data. Axes represent the summarization of variability in the data set, the greater the distance between points, the greater the dissimilarity. Sample grouping was found to explain significant variation in overall bacterial dissimilarity (PERMANOVA; *p* = 0.001, R^2^ = 0.13). An ellipse was added for each group and shows a 95% confidence interval. d) The Arcsine-transformed relative abundance of *S. aureus* and *S. epidermidis* grouped and colored by the sample groups. Both species were indicated to be differentially abundant per ANCOM-BC with pairwise testing by Tukey’s HSD test. Significance asterisks represent pairwise testing, where ‘ns’=p>0.05; ‘*’=0.01<p<0.05; ‘**’= 0.001<p<0.01; and ‘***’= p<0.001.

### Metabolomics molecular profile

Overall, the processed mass spectrometry data resulted in 9,541 MS^1^ features (*i*.*e*., a detected signal with *m/z* and retention time corresponding to a detected molecule). MS^2^ spectra were collected for all MS^1^ features, which were represented in a molecular network (FIG S1a) based on spectral similarity (21). Around 9.5% of the MS^2^ spectra were annotated by library matches against the reference GNPS public libraries, which is similar to the ∼10% annotation rate reported for other human matrices (22). Some examples of annotated molecules are illustrated in FIG S1b. It is important to note that the suspect library (23), an *in silico* MS/MS library utilized to propagate annotation on GNPS, accounted for around 80% of the annotations. We used the MS features to create pairwise PLS-DA models. FIG S2 (a,b) shows a remarkable separation between healthy vs. ADNL, as well as between healthy vs. ADL, suggesting that the metabolic profiles of both ADNL and ADL are distinct from the healthy group. For this, we employed a method called random forest classifier and trained it on the dataset using cross-validation. By measuring the decrease in classification accuracy when each feature was randomly permuted, we filtered the 15 most informative features for distinguishing between the groups (FIG 3a). Then, we annotated three out of the top five molecules that contribute to classifying different groups, such as aspartyl-phenylalanine, leucylproline, *N*-acetyl-methionine, and others more abundant in ADL samples. Five of the top 15 metabolites were not annotated, and it can be attributed to limited MS/MS databases or even to novel molecules. To visualize the results, we created box plots (FIG 3b) to visualize the distribution of the normalized peak intensities of six features across the different groups (healthy, ADNL, and ADL). The significant differences in relative concentration for these molecules per group were assessed by ANOVA analysis (*p-*value < 0.05). Aspartyl-phenylalanine, leucylproline, *N*-acetyl-methionine, phytosphingosine, and two related compounds were annotated at level 2 by GNPS libraries.

**FIG 3.**
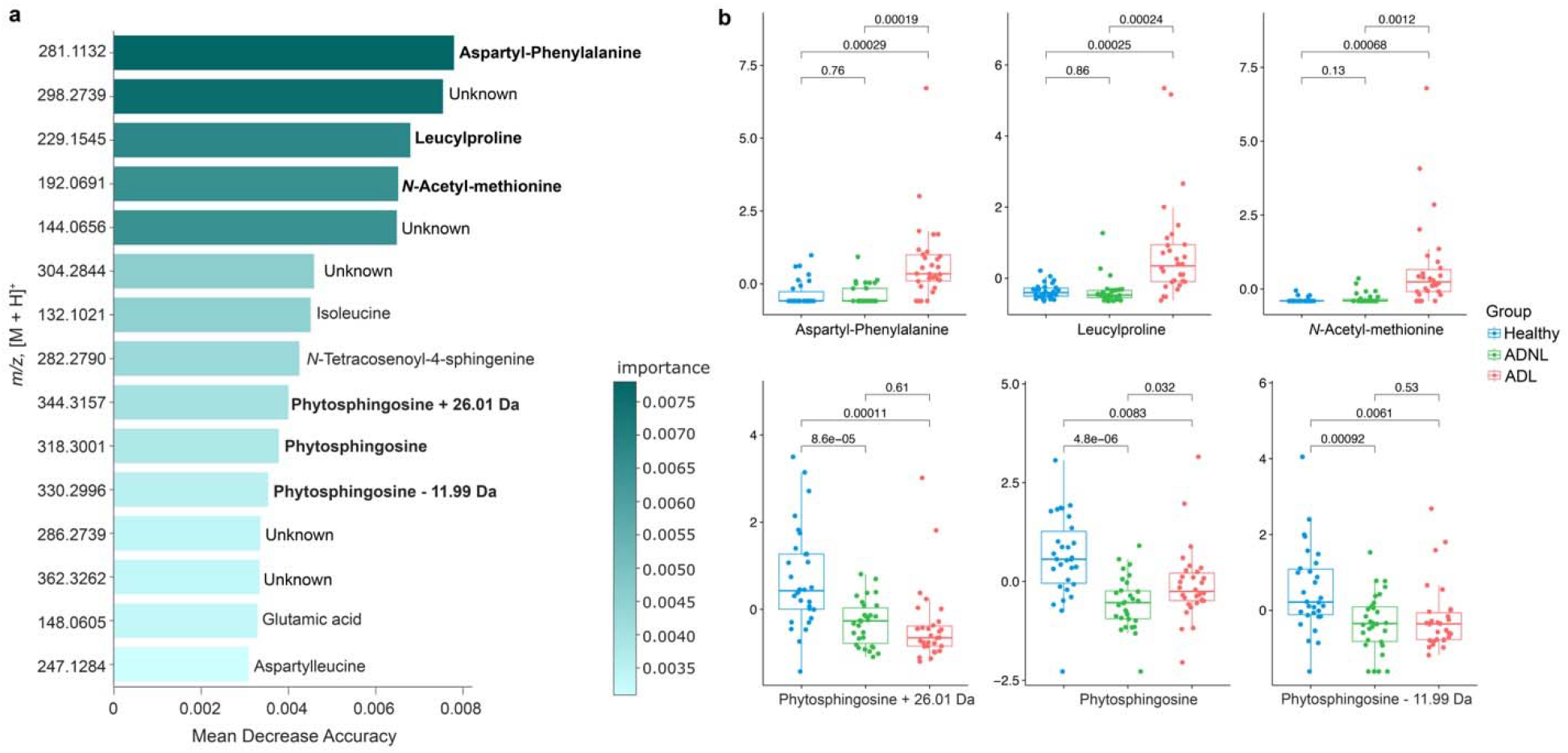
Mapping the main molecules in healthy and Atopic dermatitis skin samples. a) The barplot contains the top 15 MS features that are differentiated from three groups: healthy, ADNL, and ADL. They were classified by random forest analysis. The MS features were ranked by mean decrease accuracy. The features with annotations and which stood out with higher concentration in ADL and healthy groups (in bold) were selected to build boxplots. b) The y-axis represents the normalized peak intensity per MS feature, and the x-axis represents the boxplots and their distribution of values for a specific MS feature by sample group. Aspartyl-phenylalanine (*m/z* 281.1132), leucylproline (*m/z* 229.1545), and *N*-acetyl-methionine (*m/z* 192.0691) were detected in the unhealthy (ADL) group. In contrast, phytosphingosine (*m/z* 318.3001) and two related compounds (*m/z* 344.3157 and 330.2996) were detected in higher concentrations in the healthy group.

### Microbiome and metabolomic integration

The integration of 16S rRNA sequencing results with metabolomics data using the mmvec (24) method enabled the prediction of co-occurrence patterns between molecules, microbial species, and sample groups. High co-occurrence probabilities indicate potential relationships, such as a microbe producing a specific molecule or a molecule inducing the proliferation of a specific microbe. The biplot in FIG 4 visually represents these microbe-molecule co-occurrences. We observed higher co-occurrence probabilities in healthy and ADNL groups compared to ADL, represented by more bacterial OTUs towards healthy and ADNL groups (FIG 4a). In the ADL group, a correlation was observed with the presence of *S. aureus*, which co-occurred with specific molecules such as aspartyl-phenylalanine, leucylproline, and *N*-acetyl-methionine, previously described in higher concentration in ADL. This suggests a potential association between *S. aureus* and the production or induction of these molecules in ADL. Furthermore, microbeMASST (25) was employed to make MS/MS single searches for the molecules detected in higher abundance in the ADL group (FIG 4b). microbeMASST is a database where users can search MS/MS spectra previously detected in experimental data from bacterial, fungal, or archaeal monoculture extracts (25). In that regard, aspartyl-phenylalanine, leucylproline, and *N*-acetyl-methionine also were detected in strains of *S. aureus*, and it is a piece of evidence these molecules could be produced by *S. aureus* strains. Also, these findings are according to the findings observed in the diversity of the skin microbiome (FIG 2d), which showed a high abundance of *S. aureus*.

**FIG 4.**
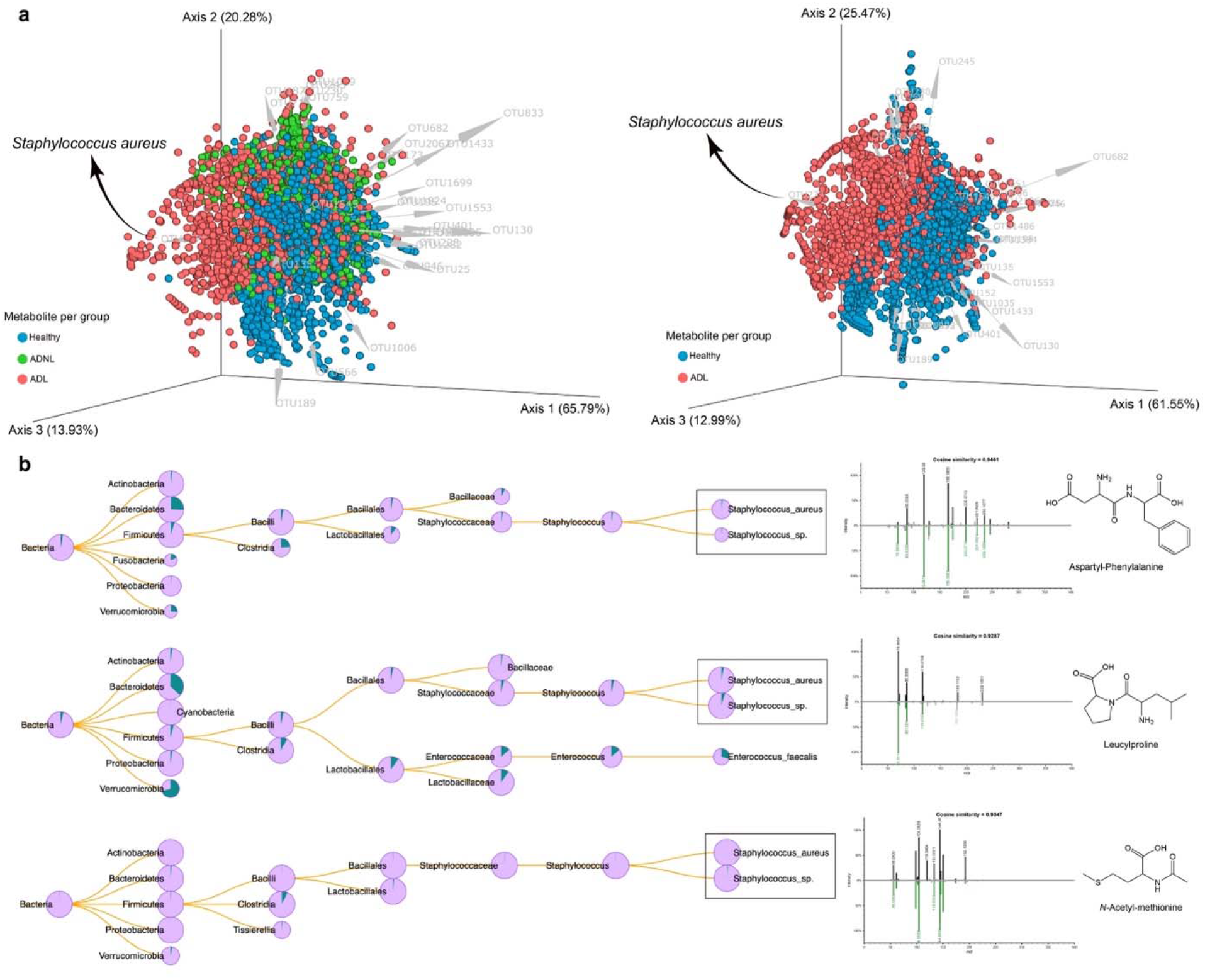
Microbe-molecule co-occurrence biplot. a) mmvec analysis included samples from the healthy, atopic dermatitis without lesions and atopic dermatitis with lesion groups (90 samples). Spheres in the biplot represent the molecules, while the arrows represent the microbes. Spheres were colored based on which group each molecule was most abundant in, while microbes were colored based on which species they belonged to. Small angles between arrows indicate microbes co-occurring with each other. Similarly, closer spheres indicate molecules co-occurring. Arrows pointing towards a group of molecules indicate microbe-molecule co-occurrence. This biplot shows the 30 most important OTUs (higher vector magnitude). b) MicrobeMASST search outputs confirm that monocultures of *S. aureus* and *S*. sp. could produce molecules such as aspartyl-phenylalanine, leucylproline, and *N*-acetyl-methionine, which were detected in higher abundance in ADL (retrieved from Random Forest analysis). Pie charts display the proportion of MS/MS matches found in the deposited reference database. Green indicates a match with the monocultures, while pink represents a nonmatch.

On the other hand, bacteria mainly present in healthy and ADNL groups such as *Paracoccus* sp, *Pseudomonas* sp, *Prevotella bivia, Lactobacillus iners, Anaerococcus* sp, *Micrococcus* sp, *Corynebacterium ureicelerivorans, Corynebacterium massiliense, Streptococcus thermophilus*, and *Roseomonas mucosa* (see FIG S3) showed associations with phytosphingosine of *m/z* 318.3001, and two related compounds of *m/z* 344.3157 and *m/z* 330.2996, respectively (FIG S1b). The 30 most important OTUs retrieved from the mmvec analysis, their taxonomy, and the molecules information (*m/z*, retention time, and spectral library annotations if available). For more details, mmvec per pairs analyses can be found in FIG S3. An alternative visualization of these co-occurrences is shown in FIGs S4-S6, in which these top 30 OTUs with co-occurrence probabilities above 5.0 are depicted (Tables S2-S4). Therefore, these results provide valuable insights into the co-occurrence patterns between microbiota and metabolites in different groups, shedding light on potential microbial-metabolite interactions and their relevance in the context of skin health.

## Discussion

In this study, to understand the underlying impacts of atopic dermatitis on human skin, we investigated the microbiome and metabolomic profiles of skin swab samples from healthy subjects and patients with atopic dermatitis. The 16S rRNA sequencing data and mass spectrometry-based metabolomics data were used to uncover the interactions between the skin microbiome and the host metabolome in atopic dermatitis. Our results demonstrated significant alterations in the skin microbiome diversity and composition in patients with atopic dermatitis. We observed a decrease in alpha diversity, as indicated by the Shannon diversity index (FIG 2a), in the atopic dermatitis lesion (ADL) group compared to both the atopic dermatitis non-lesion (ADNL) group and the healthy group. This finding suggests a dysbiosis in the skin microbiome of patients with atopic dermatitis (26, 27), particularly in those with active lesions. The reduced diversity may indicate a loss of beneficial microbial species and a shift towards a less diverse and potentially pathogenic microbiota (28).

Beta diversity analysis using weighted UniFrac distances further supported the distinct microbial composition in atopic dermatitis lesions compared to the non-lesion and healthy groups. The samples from the atopic dermatitis lesion group exhibited a clear separation from the other two groups in the principal coordinates analysis (PCoA) plot. The differential analysis of bacterial species using ANCOM-BC revealed specific taxa significantly associated with atopic dermatitis lesions (FIG 2b). *S. aureus* and *S. epidermidis* were identified as the most representative bacteria with significantly higher abundances in the atopic dermatitis lesion group (FIG 2d). These findings are consistent with previous studies that have reported the overgrowth of *Staphylococcus* species in atopic dermatitis lesions, suggesting their potential role in disease pathogenesis and exacerbation, particularly dominated by *S. aureus* strains (26, 27, 29). These results indicate that active lesions in atopic dermatitis are associated with a distinct microbiome composition, highlighting the importance of the skin microbiota in the pathogenesis of the disease. *S. epidermidis* is a well-known beneficial bacterium on the skin surface. It produces antimicrobial peptides to inhibit pathogenic bacteria, *e*.*g*., *S. aureus* (30), produces ceramides to maintain the skin barrier function (31), and regulars host immunity (32). However, we detected that *S. epidermidis* is also significantly increased in AD lesions. The increase in *S. epidermidis* could be due to a compensatory or antagonistic mechanism in an attempt to control *S. aureus* (7, 30–32).

Additionally, the skin metabolomic profiles were also different between the groups. Random forest analysis retrieved the top molecules and their contributions to the separation of the healthy, ADNL, and ADL groups. Noteworthy, the top 15 features most important for the classification were detected in high relative abundance in ADL samples. Among these molecules, three matches against GNPS spectral libraries were retrieved, *i*.*e*., aspartyl-phenylalanine, leucylproline, and *N*-acetyl-methionine. aspartyl-phenylalanine is a dipeptide metabolic byproduct of aspartame (*N*-L-α-aspartyl-L-phenylalanine 1-methyl ester). However, there is not a clear connection between consumption of aspartame and atopic dermatitis. In the same way, insights into leucylproline also are not established yet. A suitable hypothesis is that they could be coming from the degradation of proteins in the human skin and could be acting as the first line of defense against infections (33, 34). To gain more information about where these molecules are coming from, we searched its MS^2^ spectra in microbeMASST, which revealed they could be produced by cell culture of *S. aureus* and *Staphylococcus* sp. These findings align with the higher abundance of *S. aureus* observed in atopic dermatitis lesions and suggest that aspartyl-phenylalanine, leucylproline, and *N*-acetyl-methionine may originate from the skin microbiome. Thus, it indicates the influence of the skin microbiome on the metabolome when submitted to the status of atopic dermatitis.

Co-occurrence analysis between metabolomics and microbial sequencing data provided insights into possible interactions between the microbiome and human skin molecules. The results of our study shed light on dysbiosis and altered crosstalk between the microbiome and metabolites in the ADL group. The compromised skin barrier observed in individuals with ADL is known to contribute to the disruption of the microbial community. This disruption leads to a reduction in communication or crosstalk between the microbiome and metabolites, suggesting an imbalance in microbial-metabolite interactions in ADL. Thus, the decrease in crosstalk observed in our study provides further evidence of dysbiosis in subjects with ADL, highlighting the importance of microbial-metabolite interactions in skin health. Furthermore, the assignment of specific molecules exclusively present in healthy individuals and ADNL samples suggests their potential protective role in skin health. These molecules, such as phytosphingosine, and two related compounds, indicate they contribute to the maintenance of skin homeostasis and protect against atopic dermatitis progression. Phytosphingosine (PHS) is a natural lipid present in the intercellular spaces of the stratum corneum of the skin. It is the one of fundamental components to maintain the skin barrier function (35). PHS has also been described with anti-microbial and inflammatory activity (36, 37). However, more research is needed to elucidate the exact functions and mechanisms of these molecules in the context of skin health and AD.

Overall, our study provides valuable insight into changes in the skin microbiome and associated metabolomic profiles. It also identifies new therapeutic targets that may be useful for developing personalized treatments for individuals with atopic dermatitis based on their unique skin microbiome and metabolic profiles.

## Materials and Methods

### Study Design and sample collection

The clinical study was conducted by ProDERM (Schenefeld, Germany). An independent Ethics committee approved the study. All the subjects signed informed consent forms. The patients with mild to moderate atopic dermatitis (AD) (N=30) with local SCOring Atopic Dermatitis (SCORAD) of at least 4 (range 0-18) in the bend of the arm and non-atopic healthy subjects (N=30) were recruited. The subjects were not allowed to use any detergents or leave cosmetics on the bend of arms at least 3 days prior sample collection. On the sample collection day, the local SCORAD of AD patients was assessed by a dermatologist or trained physician. Then, two skin swab samples were collected from the lesion and non-lesion (5 cm away from the lesion) site in AD patients. Two swabs were also collected in the bend of the arm in healthy subjects. One swab was used for 16S rRNA gene sequencing analysis; another one was used for metabolomics analysis.

### 16S rRNA gene sequencing and data analysis

V1-3 region of 16S rRNA gene sequencing was conducted by RTL Genomics (Lubbock, Texas) (38). Paired reads were assembled, quality filtered, and clustered into 97% sequence similarity OTUs. Taxonomic classification was performed using an RTL Genomics classifier and an *in-house* reference database. The alpha and beta diversity differences between AD lesion, AD nonlesion, and the healthy group were analyzed using ANOVA and PERMANOVA, respectively. ANCOM-BC (39) procedure was used to screen differentially abundant taxa between groups. More details can be found in Supplementary Materials and Methods.

### Metabolomics and data processing

The cotton buds of all samples were added into a 96-deep well plate (2 mL). Aliquots of 500 μL of ethanol/H_2_O (1:1) were added to each well for extraction. The plates were sonicated for 5 min in an ultrasound bath (Danbury, CT, USA), vortexed (∼10 s), and then, the swabs were removed and then dried by speed vacuum. The samples were resuspended with 200 μL of ACN/H_2_O (1:1) and transferred to the 200 μL ThermoScientific 96-well plate for LC-MS/MS analysis. Expanded methods are provided in Supplementary Materials and Methods.

The metabolomic analyses were performed in a Vanquish UHPLC system coupled to a Q-Exactive Orbitrap mass spectrometer (Thermo Fisher Scientific, Waltham, MA), controlled by Thermo SII for Xcalibur software (Thermo Fisher Scientific, Waltham, MA). The chromatographic analysis was carried out on a Kinetex C18 column (50 × 2.1 mm, 1.7 μm particle size, 100 A pore size, Phenomenex, Torrance, USA). The gradient elution was carried out using ultra-pure water (solvent A) and acetonitrile (solvent B), both acidified with 0.1 % of formic acid (FA). The data were acquired in an *m/z* range from 80 to 2,000 Da with an electrospray source operating in the positive ionization mode. More details can be found in Supplementary Materials and Methods.

The LC-MS/MS data were processed using MZmine 3.1.0 (40). The outputs containing MS features were exported as a .mgf file and a .csv file, which were then used in the Feature-Based Molecular Networking workflow (21) on the GNPS platform (https://gnps.ucsd.edu/) and downstream statistical analysis. The job can be found at the following link (https://gnps.ucsd.edu/ProteoSAFe/status.jsp?task=a36305db40414b6ea9c5b9a5de380bea) and the parameters used can be found in supplementary materials and methods. Statistical analyses were performed in Pythin using Jupyter Notebook. Random Forest was performed using the Random Forest Classifier Scikit-Learn package. Differences in samples were tested with one-way ANOVA followed by the Tukey HSD test. Multivariate analysis was performed using the ‘pandas’, ‘sklearn.decomposition.PCA’, and ‘sklearn.cross_decomposition.PLSRegression’ packages. The methods detailed are available in Supplementary Materials and Methods.

### Microbiome and metabolomic correlation analysis

Co-occurrence probabilities between microbes and molecules were calculated using the mmvec (microbial-molecule vectors) tool (24). The relative abundance matrix for the sequencing data and the MS feature abundance table were used as inputs. Emperor (41) was used to inspect the feature-feature biplots visually. Co-occurrence probabilities were also visualized as networks using Cytoscape 3.9.1 (42). Expanded methods are provided in Supplementary Materials and Methods.

## Ethical statements

This study was reviewed and approved by an Ethics Committee (2021-100691-BO-ff) at Colgate, and the participants provided written informed consent.

## Data availability statement

The 16S rRNA sequencing data is publicly available at https://www.ncbi.nlm.nih.gov/bioproject/998761. The raw data of metabolomics are publicly available online at MassIVE (https://massive.ucsd.edu/) under the accession MSV000090788. All data generated or analyzed during this study are included in this published article (and its Supplementary Information Files).

## Acknowledgments

Colgate-Palmolive Company supported this work.

## Supplementary material

Supplementary Materials and Methods is available online only. FIG S1. Molecular network and examples of annotated molecules. FIG S2. Different groups lead to distinct skin biochemical profiles. FIG S3. Microbe-molecule co-occurrence biplots. FIG S4. The network was obtained from the mmvec co-occurrences probabilities for all the samples in the study (90 samples, three groups). FIG S5. The network was obtained from the mmvec co-occurrences probabilities for the samples from the healthy and atopic dermatitis without lesion groups (60 samples). FIG S6. The network was obtained from the mmvec co-occurrences probabilities for the samples from the healthy and atopic dermatitis with lesion groups (60 samples). Table S1. Species Identified by ANCOMBC vary between the groups. Table S2. Top 30 OTUs highlighted in the biplots obtained from mmvec analysis. All groups (Healthy, ADNL, and ADL). Table S3. Top 30 OTUs highlighted in the biplots obtained from mmvec analysis. The OTUS were retrieved from pair analysis between Healthy and ADNL groups. Table S4. Top 30 OTUs highlighted in the biplots obtained from mmvec analysis. The OTUS were retrieved from pair analysis between Healthy and ADL groups.

## References

1. Roux P-F, Oddos T, Stamatas G. 2022. Deciphering the Role of Skin Surface Microbiome in Skin Health: An Integrative Multiomics Approach Reveals Three Distinct Metabolite-Microbe Clusters. J Invest Dermatol 142:469–479.e5.

2. Blander JM, Longman RS, Iliev ID, Sonnenberg GF, Artis D. 2017. Regulation of inflammation by microbiota interactions with the host. Nat Immunol 18:851–860.

3. Byrd AL, Belkaid Y, Segre JA. 2018. The human skin microbiome. Nat Rev Microbiol 16:143–155.

4. Kim BE, Leung DYM. 2018. Significance of Skin Barrier Dysfunction in Atopic Dermatitis. Allergy Asthma Immunol Res 10:207–215.

5. Sharifi-Rad M, Anil Kumar NV, Zucca P, Varoni EM, Dini L, Panzarini E, Rajkovic J, Tsouh Fokou PV, Azzini E, Peluso I, Prakash Mishra A, Nigam M, El Rayess Y, Beyrouthy ME, Polito L, Iriti M, Martins N, Martorell M, Docea AO, Setzer WN, Calina D, Cho WC, Sharifi-Rad J. 2020. Lifestyle, Oxidative Stress, and Antioxidants: Back and Forth in the Pathophysiology of Chronic Diseases. Front Physiol 11:694.

6. De Pessemier B, Grine L, Debaere M, Maes A, Paetzold B, Callewaert C. 2021. Gut-Skin Axis: Current Knowledge of the Interrelationship between Microbial Dysbiosis and Skin Conditions. Microorganisms 9.

7. Kong HH, Oh J, Deming C, Conlan S, Grice EA, Beatson MA, Nomicos E, Polley EC, Komarow HD, NISC Comparative Sequence Program, Murray PR, Turner ML, Segre JA. 2012. Temporal shifts in the skin microbiome associated with disease flares and treatment in children with atopic dermatitis. Genome Res 22:850–859.

8. Nakatsuji T, Gallo RL. 2019. The role of the skin microbiome in atopic dermatitis. Ann Allergy Asthma Immunol 122:263–269.

9. Randhawa M, Southall M, Samaras ST. 2013. Metabolomic analysis of sun exposed skin. Mol Biosyst 9:2045–2050.

10. Bouslimani A, Porto C, Rath CM, Wang M, Guo Y, Gonzalez A, Berg-Lyon D, Ackermann G, Moeller Christensen GJ, Nakatsuji T, Zhang L, Borkowski AW, Meehan MJ, Dorrestein K, Gallo RL, Bandeira N, Knight R, Alexandrov T, Dorrestein PC. 2015. Molecular cartography of the human skin surface in 3D. Proc Natl Acad Sci U S A 112:E2120–9.

11. Bouslimani A, da Silva R, Kosciolek T, Janssen S, Callewaert C, Amir A, Dorrestein K, Melnik AV, Zaramela LS, Kim J-N, Humphrey G, Schwartz T, Sanders K, Brennan C, Luzzatto-Knaan T, Ackermann G, McDonald D, Zengler K, Knight R, Dorrestein PC. 2019. The impact of skin care products on skin chemistry and microbiome dynamics. BMC Biol 17:47.

12. He J, Jia Y. 2022. Application of omics technologies in dermatological research and skin management. J Cosmet Dermatol 21:451–460.

13. Elpa DP, Chiu H-Y, Wu S-P, Urban PL. 2021. Skin Metabolomics. Trends Endocrinol Metab 32:66–75.

14. Alseekh S, Aharoni A, Brotman Y, Contrepois K, D’Auria J, Ewald J, C Ewald J, Fraser PD, Giavalisco P, Hall RD, Heinemann M, Link H, Luo J, Neumann S, Nielsen J, Perez de Souza L, Saito K, Sauer U, Schroeder FC, Schuster S, Siuzdak G, Skirycz A, Sumner LW, Snyder MP, Tang H, Tohge T, Wang Y, Wen W, Wu S, Xu G, Zamboni N, Fernie AR. 2021. Mass spectrometry-based metabolomics: a guide for annotation, quantification and best reporting practices. Nat Methods 18:747–756.

15. Doerr A. 2016. Global metabolomics. Nat Methods 14:32–32.

16. Fessenden M. 2016. Metabolomics: Small molecules, single cells. Nature 540:153–155.

17. Thompson LR, Sanders JG, McDonald D, Amir A, Ladau J, Locey KJ, Prill RJ, Tripathi A, Gibbons SM, Ackermann G, Navas-Molina JA, Janssen S, Kopylova E, Vázquez-Baeza Y, González A, Morton JT, Mirarab S, Zech Xu Z, Jiang L, Haroon MF, Kanbar J, Zhu Q, Jin Song S, Kosciolek T, Bokulich NA, Lefler J, Brislawn CJ, Humphrey G, Owens SM, Hampton-Marcell J, Berg-Lyons D, McKenzie V, Fierer N, Fuhrman JA, Clauset A, Stevens RL, Shade A, Pollard KS, Goodwin KD, Jansson JK, Gilbert JA, Knight R, Earth Microbiome Project Consortium. 2017. A communal catalogue reveals Earth’s multiscale microbial diversity. Nature 551:457–463.

18. Ziemert N, Alanjary M, Weber T. 2016. The evolution of genome mining in microbes - a review. Nat Prod Rep 33:988–1005.

19. Shaffer JP, Nothias L-F, Thompson LR, Sanders JG, Salido RA, Couvillion SP, Brejnrod AD, Lejzerowicz F, Haiminen N, Huang S, Lutz HL, Zhu Q, Martino C, Morton JT, Karthikeyan S, Nothias-Esposito M, Dührkop K, Böcker S, Kim HW, Aksenov AA, Bittremieux W, Minich JJ, Marotz C, Bryant MM, Sanders K, Schwartz T, Humphrey G, Vásquez-Baeza Y, Tripathi A, Parida L, Carrieri AP, Beck KL, Das P, González A, McDonald D, Ladau J, Karst SM, Albertsen M, Ackermann G, DeReus J, Thomas T, Petras D, Shade A, Stegen J, Song SJ, Metz TO, Swafford AD, Dorrestein PC, Jansson JK, Gilbert JA, Knight R, Earth Microbiome Project 500 (EMP500) Consortium. 2022. Standardized multi-omics of Earth’s microbiomes reveals microbial and metabolite diversity. Nat Microbiol 7:2128–2150.

20. Peng W, Huang J, Yang J, Zhang Z, Yu R, Fayyaz S, Zhang S, Qin Y-H. 2019. Integrated 16S rRNA Sequencing, Metagenomics, and Metabolomics to Characterize Gut Microbial Composition, Function, and Fecal Metabolic Phenotype in Non-obese Type 2 Diabetic Goto-Kakizaki Rats. Front Microbiol 10:3141.

21. Nothias L-F, Petras D, Schmid R, Dührkop K, Rainer J, Sarvepalli A, Protsyuk I, Ernst M, Tsugawa H, Fleischauer M, Aicheler F, Aksenov AA, Alka O, Allard P-M, Barsch A, Cachet X, Caraballo-Rodriguez AM, Da Silva RR, Dang T, Garg N, Gauglitz JM, Gurevich A, Isaac G, Jarmusch AK, Kameník Z, Kang KB, Kessler N, Koester I, Korf A, Le Gouellec A, Ludwig M, Martin H C, McCall L-I, McSayles J, Meyer SW, Mohimani H, Morsy M, Moyne O, Neumann S, Neuweger H, Nguyen NH, Nothias-Esposito M, Paolini J, Phelan VV, Pluskal T, Quinn RA, Rogers S, Shrestha B, Tripathi A, van der Hooft JJJ, Vargas F, Weldon KC, Witting M, Yang H, Zhang Z, Zubeil F, Kohlbacher O, Böcker S, Alexandrov T, Bandeira N, Wang M, Dorrestein PC. 2020. Feature-based molecular networking in the GNPS analysis environment. Nat Methods 17:905–908.

22. Gomes PWP, Zuffa S, Baumeister A, Caraballo-Rodríguez AM, Zhao HN, Mannochio-Russo H, North M, Dogo-isonagie C, Patel O, Lavender S, Pimenta P, Gronlund J, Pilch S, Maloney V, Dorrestein PC. 2023. The effects of bleaching strategies on the teeth metabolome.

23. Bittremieux W, Avalon NE, Thomas SP, Kakhkhorov SA, Aksenov AA, Gomes PWP, Aceves CM, Rodríguez AMC, Gauglitz JM, Gerwick WH, Jarmusch AK, Kaddurah-Daouk RF, Kang KB, Kim HW, Kondić T, Mannochio-Russo H, Meehan MJ, Melnik AV, Nothias L-F, O’Donovan C, Panitchpakdi M, Petras D, Schmid R, Schymanski EL, van der Hooft JJJ, Weldon KC, Yang H, Zemlin J, Wang M, Dorrestein PC. 2022. Open Access Repository-Scale Propagated Nearest Neighbor Suspect Spectral Library for Untargeted Metabolomics. bioRxiv.

24. Morton JT, Aksenov AA, Nothias LF, Foulds JR, Quinn RA, Badri MH, Swenson TL, Van Goethem MW, Northen TR, Vazquez-Baeza Y, Wang M, Bokulich NA, Watters A, Song SJ, Bonneau R, Dorrestein PC, Knight R. 2019. Learning representations of microbe-metabolite interactions. Nat Methods 16:1306–1314.

25. Zuffa S, Schmid R, Bauermeister A, Gomes PWP, Caraballo-Rodriguez AM, El Abiead Y, Aron AT, Gentry EC, Zemlin J, Meehan MJ, Avalon NE, Cichewicz RH, Buzun E, Terrazas MC, Hsu C-Y, Oles R, Ayala AV, Zhao J, Chu H, Kuijpers MCM, Jackrel SL, Tugizimana F, Nephali LP, Dubery IA, Madala NE, Moreira EA, Costa-Lotufo LV, Lopes NP, Rezende-Teixeira P, Jimenez PC, Rimal B, Patterson AD, Traxler MF, de Cassia Pessotti R, Alvarado-Villalobos D, Tamayo-Castillo G, Chaverri P, Escudero-Leyva E, Quiros-Guerrero L-M, Bory AJ, Joubert J, Rutz A, Wolfender J-L, Allard P-M, Sichert A, Pontrelli S, Pullman BS, Bandeira N, Gerwick WH, Gindro K, Massana-Codina J, Wagner BC, Forchhammer K, Petras D, Aiosa N, Garg N, Liebeke M, Bourceau P, Kang KB, Gadhavi H, de Carvalho LPS, dos Santos MS, Perez-Lorente AI, Molina-Santiago C, Romero D, Franke R, Bronstrup M, de Leon AVP, Pope PB, La Rosa SL, La Barbera G, Roager HM, Laursen MF, Hammerle F, Siewert B, Peintner U, Licona-Cassani C, Rodriguez-Orduna L, Rampler E, Hildebrand F, Koellensperger G, Schoeny H, Hohenwallner K, Panzenboeck L, Gregor R, O’Neill EC, Roxborough ET, Odoi J, Bale NJ, Ding S, Sinninghe Damste JS, Guan XL, Cui JJ, Ju K-S, Silva DB, Silva FMR, da Silva GF, Koolen HHF, Grundmann C, Clement JA, Mohimani H, Broders K, McPhail KL, Ober-Singleton SE, Rath CM, McDonald D, Knight R, Wang M, Dorrestein PC. 2023. A Taxonomically-informed Mass Spectrometry Search Tool for Microbial Metabolomics Data. bioRxiv.

26. Khadka VD, Key FM, Romo-González C, Martínez-Gayosso A, Campos-Cabrera BL, Gerónimo-Gallegos A, Lynn TC, Durán-McKinster C, Coria-Jiménez R, Lieberman TD, García-Romero MT. 2021. The Skin Microbiome of Patients With Atopic Dermatitis Normalizes Gradually During Treatment. Front Cell Infect Microbiol 11:720674.

27. Kobayashi T, Glatz M, Horiuchi K, Kawasaki H, Akiyama H, Kaplan DH, Kong HH, Amagai M, Nagao K. 2015. Dysbiosis and Staphylococcus aureus Colonization Drives Inflammation in Atopic Dermatitis. Immunity 42:756–766.

28. DeGruttola AK, Low D, Mizoguchi A, Mizoguchi E. 2016. Current Understanding of Dysbiosis in Disease in Human and Animal Models. Inflamm Bowel Dis 22:1137–1150.

29. Ogonowska P, Gilaberte Y, Barańska-Rybak W, Nakonieczna J. 2020. Colonization With Staphylococcus aureus in Atopic Dermatitis Patients: Attempts to Reveal the Unknown. Front Microbiol 11:567090.

30. Cogen AL, Yamasaki K, Muto J, Sanchez KM, Crotty Alexander L, Tanios J, Lai Y, Kim JE, Nizet V, Gallo RL. 2010. Staphylococcus epidermidis antimicrobial delta-toxin (phenol-soluble modulin-gamma) cooperates with host antimicrobial peptides to kill group A Streptococcus. PLoS One 5:e8557.

31. Zheng Y, Hunt RL, Villaruz AE, Fisher EL, Liu R, Liu Q, Cheung GYC, Li M, Otto M. 2022. Commensal Staphylococcus epidermidis contributes to skin barrier homeostasis by generating protective ceramides. Cell Host Microbe 30:301–313.e9.

32. Gallo RL. 2015. S. epidermidis influence on host immunity: more than skin deep. Cell Host Microbe.

33. Graf M, Wilson DN. 2019. Intracellular Antimicrobial Peptides Targeting the Protein Synthesis Machinery. Adv Exp Med Biol 1117:73–89.

34. Patriarca EJ, Cermola F, D’Aniello C, Fico A, Guardiola O, De Cesare D, Minchiotti G. 2021. The Multifaceted Roles of Proline in Cell Behavior. Front Cell Dev Biol 9:728576.

35. Knox S, O’Boyle NM. 2021. Skin lipids in health and disease: A review. Chem Phys Lipids 236:105055.

36. Pavicic T, Wollenweber U, Farwick M, Korting HC. 2007. Anti-microbial and -inflammatory activity and efficacy of phytosphingosine: an in vitro and in vivo study addressing acne vulgaris. Int J Cosmet Sci 29:181–190.

37. Rollin-Pinheiro R, Singh A, Barreto-Bergter E, Del Poeta M. 2016. Sphingolipids as targets for treatment of fungal infections. Future Med Chem 8:1469–1484.

38. Tipton CD, Sanford NE, Everett JA, Gabrilska RA, Wolcott RD, Rumbaugh KP, Phillips CD. 2019. Chronic wound microbiome colonization on mouse model following cryogenic preservation. PLoS One 14:e0221565.

39. Lin H, Peddada SD. 2020. Analysis of compositions of microbiomes with bias correction. Nat Commun 11:3514.

40. Schmid R, Heuckeroth S, Korf A, Smirnov A, Myers O, Dyrlund TS, Bushuiev R, Murray KJ, Hoffmann N, Lu M, Sarvepalli A, Zhang Z, Fleischauer M, Dührkop K, Wesner M, Hoogstra SJ, Rudt E, Mokshyna O, Brungs C, Ponomarov K, Mutabdžija L, Damiani T, Pudney CJ, Earll M, Helmer PO, Fallon TR, Schulze T, Rivas-Ubach A, Bilbao A, Richter H, Nothias L-F, Wang M, Orešič M, Weng J-K, Böcker S, Jeibmann A, Hayen H, Karst U, Dorrestein PC, Petras D, Du X, Pluskal T. 2023. Integrative analysis of multimodal mass spectrometry data in MZmine 3. Nat Biotechnol 41:447–449.

41. Vázquez-Baeza Y, Pirrung M, Gonzalez A, Knight R. 2013. EMPeror: a tool for visualizing high-throughput microbial community data. GigaScience https://doi.org/10.1186/2047-217x-2-16.

42. Shannon P, Markiel A, Ozier O, Baliga NS, Wang JT, Ramage D, Amin N, Schwikowski B, Ideker T. 2003. Cytoscape: a software environment for integrated models of biomolecular interaction networks. Genome Res 13:2498–2504.

